# RGEN-SEQ FOR HIGHLY SENSITIVE AMPLIFICATION-FREE SCREEN OF OFF-TARGET SITES OF GENE EDITORS

**DOI:** 10.1101/2021.07.01.450795

**Authors:** Alexander Kuzin, Brendan Redler, Jaya Onuska, Alexei Slesarev

## Abstract

Sensitive detection of off-target sites produced by gene editing nucleases is crucial for developing reliable gene therapy platforms. Although several biochemical assays for the characterization of nuclease off-target effects have been recently published, they still leave plenty of room for improvement. Here we describe a sensitive, PCR-free next-generation sequencing method (RGEN-seq) for unbiased detection of double-stranded breaks generated by RNA-guided CRISPR-Cas9 endonuclease. The method is extremely simple, and it is on a par or even supersedes in sensitivity existing assays without reliance on amplification steps. The latter saves time, simplifies workflow, and removes genomic coverage bias and gaps associated with PCR and/or other enrichment procedures. RGEN-seq is fully compatible with existing off-target detection software; moreover, the unbiased nature of RGEN-seq offers a robust foundation for relating assigned DNA cleavage scores to propensity for off-target mutations in cells. A detailed comparison of RGEN-seq with other off-target detection methods is provided using a previously characterized set of guide RNAs.

## Introduction

The CRISPR–Cas9 system is a potent genome editing tool for biology^1–3^, and has a great potential for gene therapy^4–6^. RNA-guided Cas9 endonuclease complexes (RGENs) became a third wave of programmable endonucleases for targeted genome editing, along with previously known zinc-finger nucleases (ZFNs)^7^ and transcription activator–like effector nucleases (TALENs)^7,8^. The rapid pace with which RGENs have been adopted for gene editing, is due to their perceived simplicity compared to other platforms. Nevertheless, high-efficiency genome editing by all platforms is based on the ability to make a targeted DNA double-strand break (DSB) in the chromosomal sequence of interest, which can be repaired by nonhomologous end-joining (NHEJ)^9^ or homology-directed repair (HDR)^10^.

One continuing concern for recruiting RGENs, ZFN, or TALENs for genome engineering is the potential for off-target DSB activity at non-consensus sites within the genome^11–13^. This concern drives developing strategies for improving targeted nuclease specificities^14–17^ as well as methods for genome-wide off-target sites detection. Cell-based and *in vivo* genome-wide methods^18–22^ rely on cellular events for the integration of donor sequences or chromosomal translocations. These methods are assumed to be precise, with few or no false positives as they operate in the context of intracellular chromatin state and modifications. However, they may not be sensitive enough to detect rare deleterious events, being confounded by site-and cell-line-dependent differences in DNA repair as well as by interactions between editing events and cell division^12,18^. By contrast, biochemical off-target assays such as DIGENOME-seq^23–25^, and in particular CIRCLE-seq^26^, CHANGE-seq^27^, and SITE-seq^28^, are exquisitely sensitive. Even though they can be prone to extensive false positives, it is becoming increasingly imperative to employ these methods to experimentally define the genome-wide activity of editing nucleases and analyze identified off-targets alongside with data obtained in cellular or *in vivo* assays.

Here we present RGEN-seq, an *in vitro* method for the genome-wide analysis of DSBs produced by genome editing nucleases, which is as simple as DIGENOME-seq and is as sensitive as SITE-seq or CIRCLE and CHANGE-seq, but which doesn’t require PCR or any other amplification steps to achieve high sensitivity and selectivity. The rationale for such a method is the fact that existing sensitive assays are multi-step and labor-intensive and include one or more amplification steps, which are a potential source of biases – read duplication, under-or over-representation of certain genomic regions^29^, depending on amplification methods used. Amplification-free methods would be easier to practice, scaleup or automate, and RGEN-seq is a step forward in this direction.

## RESULTS

### RGEN-seq method

RGEN-seq relies on Illumina sequencing workflow and is similar to CIRCLE/CHANGE-seq and SITE-seq in that it is based on tagging RGEN cut sites and selectively sequencing cleaved genomic DNA but it employs fewer biochemical steps and avoids the use of DNA circularization, PCR, or affinity capture enrichment (Fig.1a; Supplementary Protocol). The high sensitivity in RGEN-seq is achieved by ligating cut ends with a truncated Y-type adaptor that lacks the p5 flowcell binding site (p7L-adapter). After a fragmentation step, the matching p5L adapter is ligated to remaining fragment ends. As a result, DNA strands containing ligated adapter oligonucleotides with both p5 and p7 sequences at their ends hybridize to the flowcell surface and generate clusters. All other DNA fragments, either without p5 and p7 sites (no ligation) or only with the p5 sequence on both fragment ends (those with two naked ends after fragmentation), cannot hybridize to the flowcell and are washed away during the clusterization process. Once sequencing is complete, reverse (R2) read sequences are mapped to a corresponding reference genome (Fig.1a). The use of R1 reads is optional, and generally we perform only 26 R1 sequencing cycles required for generating clusters. An alternative RGEN-seq variant, which requires only R1 reads can be used too (Supplementary Fig. s1); though in this case non-productive adapters can anneal to flowcell oligos and potentially reduce the sequencing run output in case of a pool of dozens of libraries.

**Figure 1.**
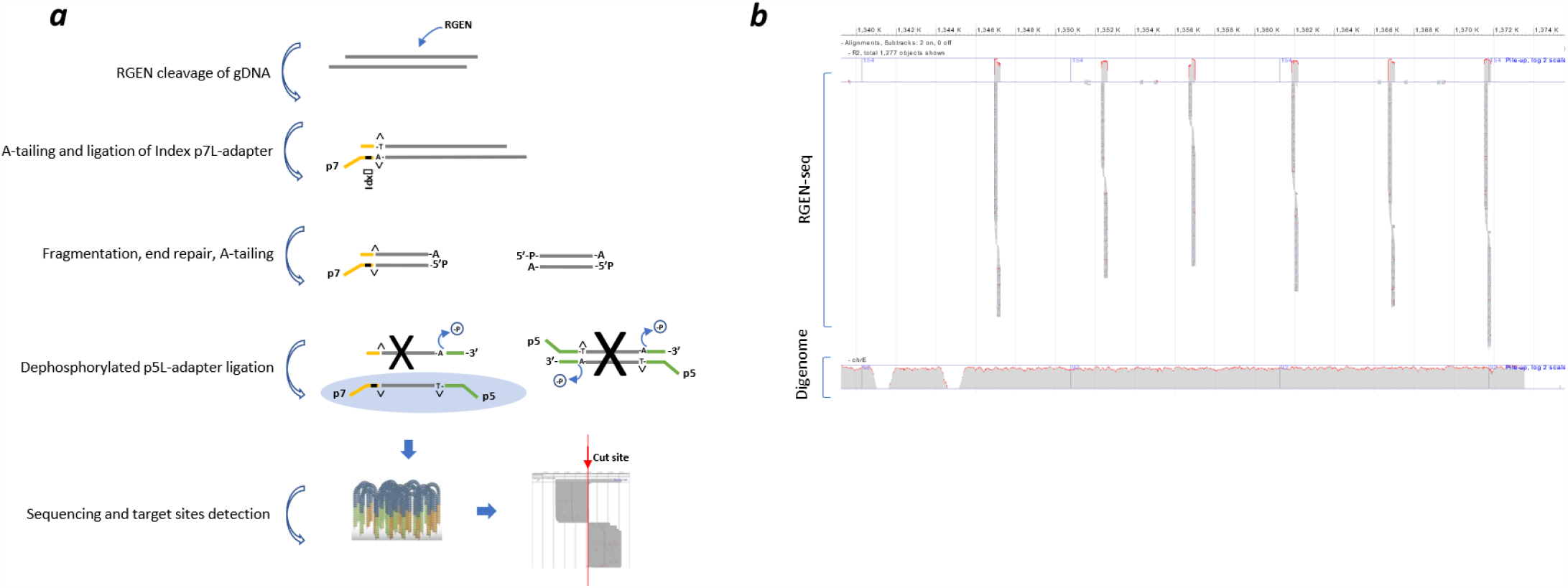
Overview of RGEN-seq workflow. (***a***), Schematic workflow illustrating preparation of the RGEN-seq library. The p7L-adapter, which is an Illumina Y-type index adapter (orange), is missing the p5 site; the p5L-adapter (green), which is a truncated Illumina Y-adapter, lacks the p7 site. Ligation of the latter adapter to both sides of free fragmented DNA prevents “background” library molecules from binding to the flowcell and generating clusters. Target and off-targets cut sites are detected by spotting the characteristic read alignment pattern around cleavage sites. (***b***), Representative IGV image of alignment around cut sites obtained using RGEN-seq with pooled six RGENs targeting an *E. coli* insertion sequence in the CHO-K1 genome (*top panel*); *bottom panel*, for comparison, read alignment in the corresponding CHO-K1 genomic region obtained using DIGENOME-seq.

In the resulting alignment, sites cleaved by the RGENs yield sequence read pileups that terminate at the cut site, ∼3 nucleotides proximal to the PAM, producing a distinct signature that can be detected computationally. This signature is similar to those observed with DIGENOME-Seq^23^, SITE-Seq^28^, and DISCOVER-Seq^22^, which allowed us to use previously developed software for the analysis of RGEN-seq experimental data^22,25,28^.

RGEN-Seq was extensively tested on a Chinese hamster CHO-K1 clone bearing a 180 kb *E. coli* insert^30^. Isolated genomic DNA was targeted with seven sgRNAs, six of which tile across a 40 kb portion of the integrated *E. coli* region (Fig. 1b), and one targeting an integrated vector sequence. Cas9 cut sites were identified with BLENDER^22^ pipeline. Figure 1b shows an IGV snapshot of the read coverage within the aforementioned genomic region -note the absence of reads between target cut sites. For comparison, about 40X whole genome coverage is required to reliably detect the sgRNA target sites in the same CHO-K1 clone using DIGENOME-seq (Fig.1b, *bottom panel*). With the most sensitive library preparation protocol, we have identified ∼100-450 cleavage sites per sgRNA, per million reads. Depending on a given cut site, RGEN-seq results in several hundred to several thousand-fold higher sensitivity than DIGENOME-seq; a result which is on par with other established *in vitro* methods.

### RGEN-seq optimization

Although RGEN-seq was designed as an amplification-free method, we didn’t want to compromise on its sensitivity and high-throughput capability. Therefore, we performed a great deal of protocol optimization in order to reduce the background read coverage and to enable the identification of off-target sites across a broad range of site cleavage sensitivity *in vitro*. A total of one hundred and fifty CHO-K1 and HEK293 RGEN-seq libraries using twenty sgRNAs were prepared testing different aspects of RGEN-seq protocol and sequenced on several Illumina sequencers (iSeq100, NextSeq 550, and NextSeq 2000). A list of applied laboratory reagents and biochemical steps (library components) used during the protocol optimization are listed in Supplementary Table s1. Ideally, in an RGEN-seq library, only genomic fragments terminated by genome editor cuts would generate clusters on the flowcell. In reality, noise is introduced from several sources as there are random background reads due to incomplete elimination of p7L-adapter, the presence of pre-existing non-specific DSBs or due to newly generated ones over the course of the procedure. Indeed, Figure 2a shows that blocking 3’ends of pre-existing single-and double-stranded breaks in the genomic template with terminal transferase and ddATP together with dephosphorylation of 5’ ends increase the percentage of the productive reads, i.e., those aligned to the RGEN-generated DNA ends. However, reducing the number of cleanup steps in the protocol has an even greater impact on the percentage of the productive reads (Fig. 2a) and so end-blocking as a pre-treatment was abandoned. Figure 2 also demonstrates the beneficial effect of end repair (LKFexo-+ dNTPs) of the Cas9-generated breaks before A-tailing compared to A-tailing alone. It is widely believed based on early observations made with plasmids and oligonucleotides that Cas9 predominantly generates blunt ends at a position three base pairs upstream of the PAM^31,32^. Given this assumption, one would expect on average an even read coverage on both sides of the cut. However, recent genome-wide screening of Cas9-generated breaks by PacBio’s and ONT’s nanopore real time sequencing^33^ methods revealed highly asymmetrical read coverage of the PAM proximal and distal regions. Figure 2b shows the coverage of the PAM proximal and distal regions is skewed in RGEN-seq too but can be leveled if the end repair (ER) step is done along with A-tailing. Together these data suggest blunt fragments are not the majority (or at least not always the majority^27^) of all ends in the Cas9 reaction and the actual structure of Cas9-generated breaks is more in line with temporal bidirectional degradation of Cas9 cleavage products, resulting in heterogeneous ends following initial DNA cleavage^34,35^.

**Figure 2.**
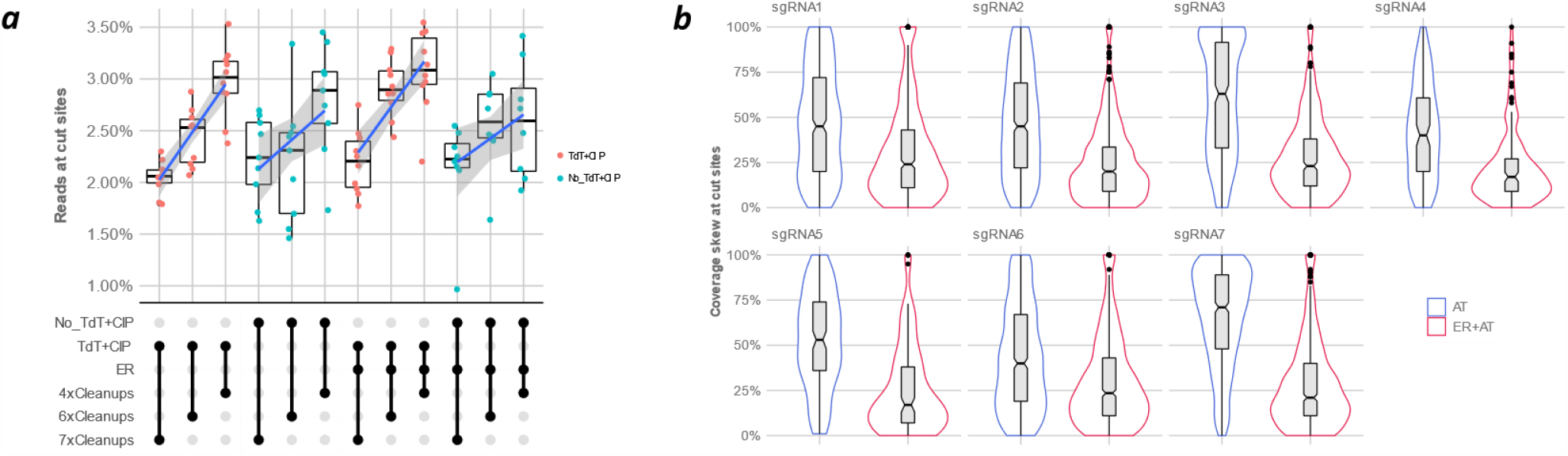
Optimization of RGEN-seq: effect of end repair. (***a***), UpSet^39^ plot showing effects of different treatment combinations on the output of productive reads as percentage of total mapped reads. *TdT*, terminal transferase; *CIP*, calf intestinal phosphatase; *ER*, DNA end repair after RGEN cleavage; *nxCleanups*, number of clean-up steps in a protocol variant (see Supplementary Protocol for details). (***b***), Leveraging skewed read coverage of the proximal and distal sequences of RGEN clevage sites. Violin plots demonstrate the effect of end repair of the cleaved sites. Coverage skewness is the normalized difference in read coverage of the proximal and distal parts of the cut site. *ER*, end repair; *AT*-A-tailing.

RGEN-seq demonstrates strong correlation of cut site scores between experiments. We performed independent library preparations with 6 guide RNAs targeted to *E. coli* insert sequence in the CHOK1 clone AD49ZG^30^ and found that BLENDER^22^ score of cut sites (roughly equal to the number of reads starting around a cut site) correlate with R^2^ > 0.92 (Fig. 3*a*). Figure 3*b* shows that although the number of recovered cut sites in CHO-K1 DNA is almost a linear function of sequencing reads used for the analysis, all target sites and off-targets with up to three mismatches (up to four mismatches in case of sgRNA6 and sgRNA7) can be recovered with just about 1 million reads. Figure 3c demonstrates the cleavage sensitivity of RGEN-seq and presents RGEN concentration-dependent recovery of target and off-target sites in HEK293 DNA using previously characterized sgRNAs^18,23–26,28^. The curves thus obtained resemble a sigmoid growth function with the logarithmic accumulation of cut sites in the ∼32-256 nM RGEN concentration range, mirroring results of SITE-seq. Off-target sites with three and more mismatches contribute to the growing number of recovered sites, while in most cases cleavage sites with 0-2 mismatches are recovered at the lowest RGEN concentration used. It is interesting to note that the BLENDER score demonstrates a different dynamic (Fig. 3*d*) – the median score of putative cut sites tends to decrease after initial increase as the RGEN concentration increases for all cut sites regardless of the numbers of mismatches. This tendency is not RGEN-seq specific because it is observed in the SITE-seq method too (Supplementary Fig. s2). Such score dynamics might appear counterintuitive, but still can be explained by considering that *S. pyogenes* Cas9 is a single-turnover enzyme^36^, which initially finds high affinity genomic sites and cleaves them efficiently without apparent dissociation^37,38^. As such sites are occupied irreversibly by Cas9, and upon further increase of Cas9 concentration, multiple low affinity sites with a high number of mismatches and corresponding low cleavage efficiency contribute to the pool of cut sites, resulting in an overall decrease of cleavage score median (Fig. 3*c,d*). More details on the protocol optimization, such as effects of p7 duplex adapter versus p7L adapter, mechanical DNA shearing versus enzymatic fragmentation, can be found in Supplementary Information.

**Figure 3.**
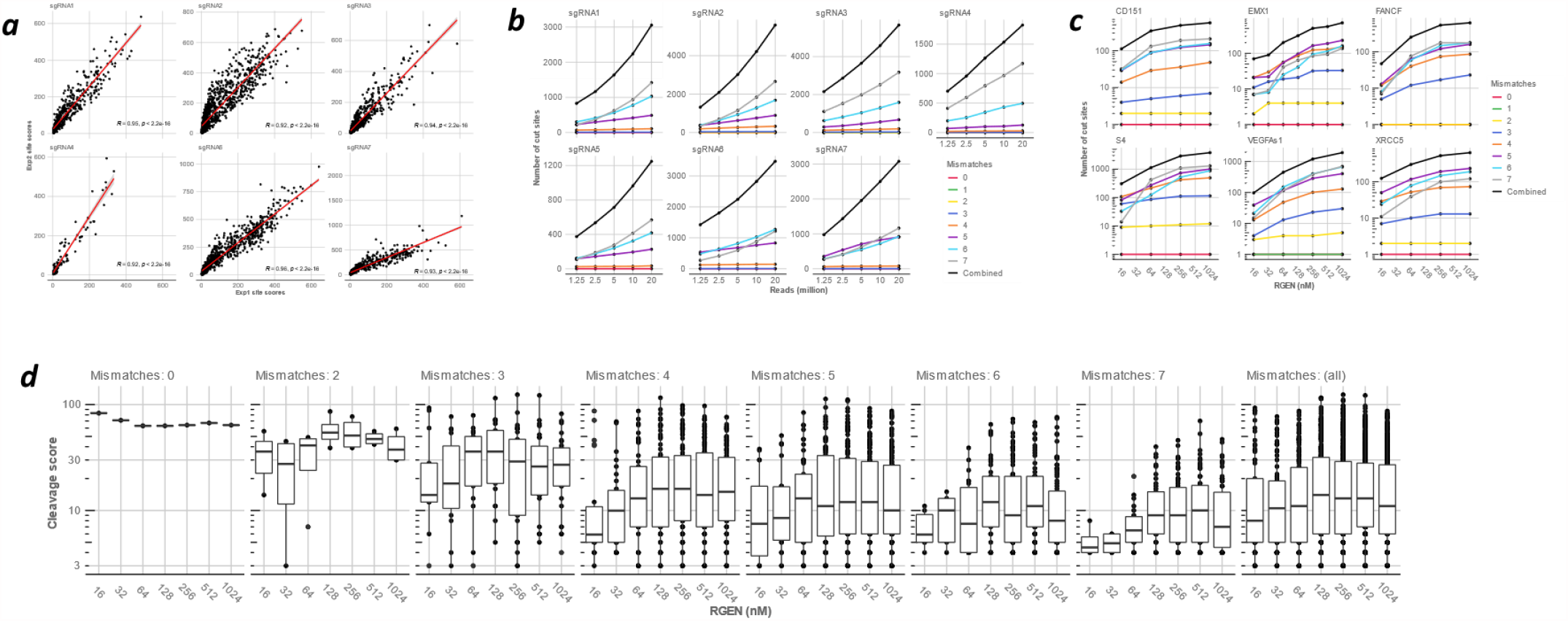
Optimization of RGEN-seq: effect of number of reads and RGEN concentration on cleavage sites recovery. (***a***) Scatterplots of RGEN-seq off-target scores between two independent libraries prepared from the same source of genomic DNA using six multiplexed sgRNAs. A Chinese hamster CHO-K1 clone bearing a 180 kb *E. coli* insert (clone AD49ZG^30^) was the source of gDNA; sgRNAs were designed to target the 180 kb *E. coli* insert sequence. Cleavage sites were called with BLENDER^22^ using two million reads unless specified otherwise. (***b***), The number of recovered RGEN-seq cut sites for seven RGENs is directly proportional to the number of total mapped reads. Genomic DNA was the same as in (***a***). (***c***), Effect of RGEN concentrations on the number of recovered off-targets using HEK293 gDNA and sgRNAs previously characterized by other off-target detection methods. (***d***), Boxplot representation of the cleavage sites score distributions as a function of the EMX1 RGEN concentration.

### Comparison of RGEN-seq with other *in vitro* off-target methods

To benchmark RGEN-seq against other biochemical genome-wide off-target detection methods, we compiled and re-analyzed data for eight different sgRNAs targeted to human genome sequences that had been previously characterized by DIGENOME-seq, CIRCLE-seq, CHANGE-seq, and SITE-seq across different human cell lines^23,25–28^. Of those eight sgRNAs, the two targeting the VEGFA site and the FANCF site had been tested by all five methods. The UpSet plot^39,40^ in Figure 4 shows shared and exclusive off-target sites for all method combinations. Sequencing read datasets from original publications were used to compare CIRCLE-seq, CHANGE-seq, and SITE-seq. To directly compare the performance of the methods, we sampled sequencing reads to ∼2 million to match the RGEN-seq sequencing depth and used BLENDER^22^ to identify off-target cut sites; for DIGENOME, only originally identified off-target sites were used^25^. Figure 4A demonstrates that in terms of the number of identified off-target sites with FANCF-specific sgRNA, RGEN-seq is on a par with SITE-seq (211 and 228 sites, respectively), while CHANGE-seq is somewhat behind with 164 sites, and CIRCLE-seq with DIGENOME producing by far the fewest sites (75 and 46). It is interesting that CHANGE-seq detects the highest number of off-target sites if used with promiscuous sgRNAs targeting VEGFAs1, S4, and EMX1 sites (Fig. 5*a,b*). In the case of more specific sgRNAs, like S1, S2, RNF2, CD151, and XRCC5, RGEN-seq supersedes other methods in terms of the number of determined off-targets (Fig. 4*b,c*).

**Figure 4.**
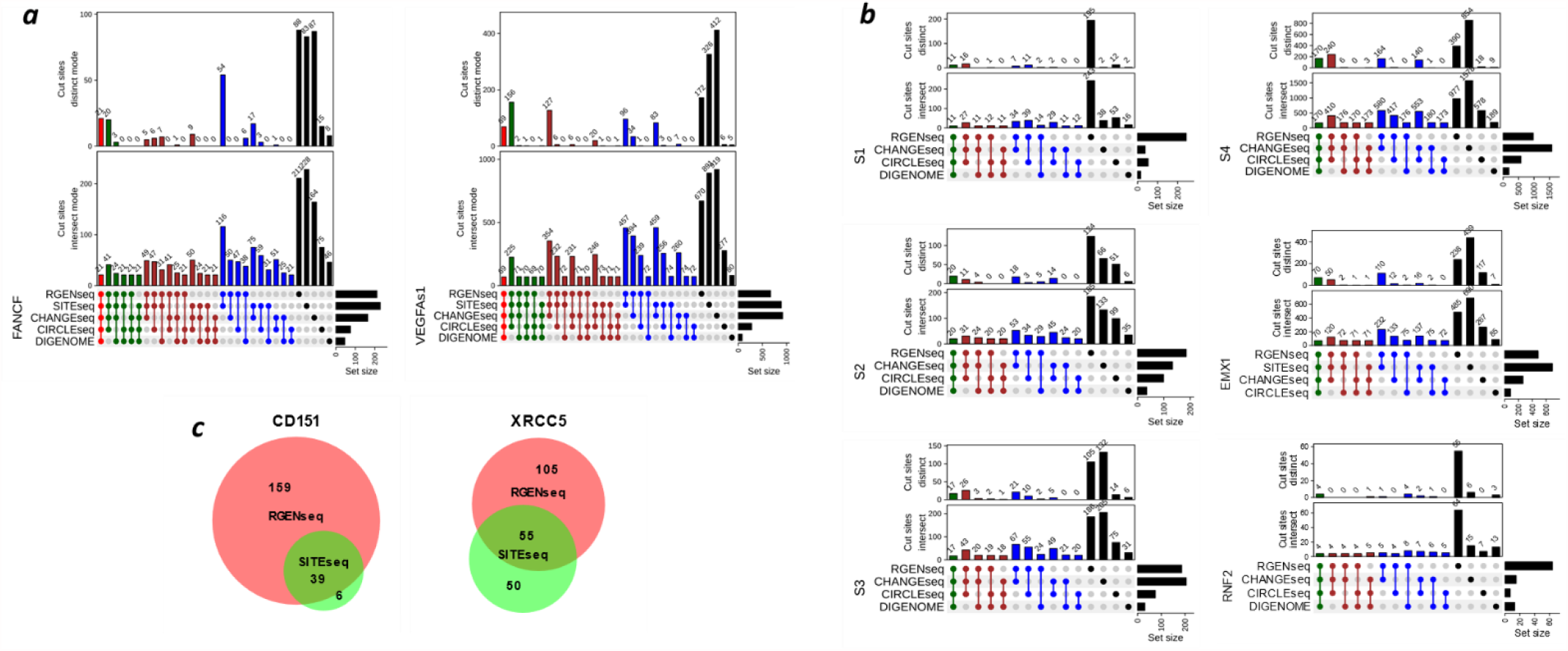
Comparison of intersecting and distinct Cas9 cut sites across different biochemical methods. (***a***) UpSet plots of intersecting (lower plot) and distinct (upper plot) off-target sites of sgRNAs against FANCF and VEGFA genes produced by five methods, and (***b***) UpSet plots of off-target sites of six sgRNAs detected by four methods. A distinct mode corresponds to exclusive intersections that contain the elements of the sets represented by the colored circles, but not of the others. Method combinations shown at the bottom of the plots are represented by solid circles colored according to the number of selected methods in the combination. (***c***) Venn diagrams showing intersecting off-target sites produced by RGEN-seq and SITE-seq using sgRNAs targeted against CD151 and XRCC5 genes. In all methods except DIGENOME, 2 million reads were used for analysis and off-target sites were identified using BLENDER^22^ with default parameters. In case of SITE-seq, CIRCLE-seq, and CHANGE-seq sequencing reads were downloaded from the NCBI’s SRA as reported in the original publications^26–28^ and analyzed in the same way as RGEN-seq data. Genomic DNA in (***a***) used in CIRCLE-seq and CHANGE-seq was from K562 and U2OS cell lines, respectively, while in RGEN-seq, SITE-seq, and DIGENOME it was from HEK293 cells. In (***b***) genomic DNA was from HEK293 cell lines in all methods in S1-S4 panels, in RGEN-seq and SITE-seq in the EMX1 panel, and in DIGENOME in the RNF2 panel. In CHANGE-seq in EMX1 and RNF2 panels DNA was from U2OS cells, in RGEN-seq and CIRCLE-seq in the RNF2 panel DNA was from K562 cells. In (***c***) DNA was from HEK293 cells.

**Figure 5.**
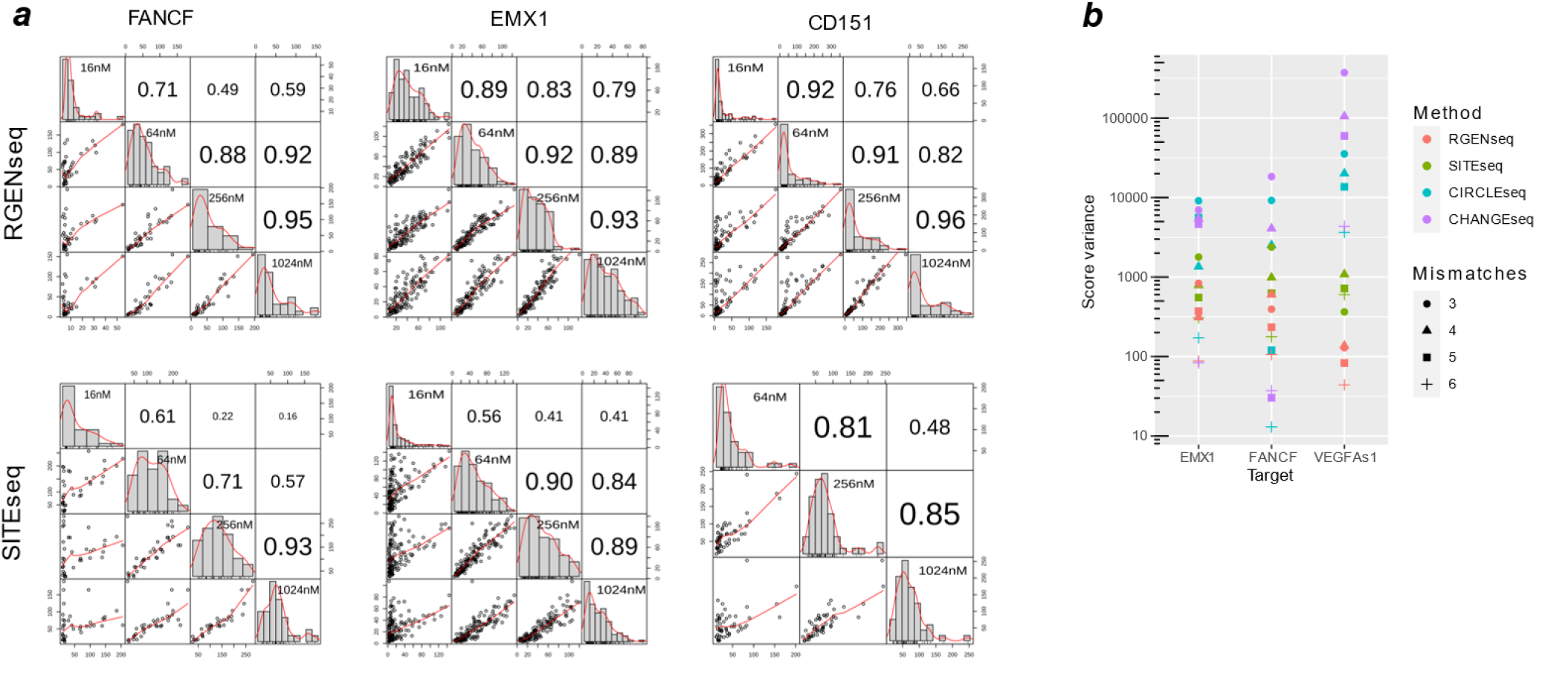
Off-target score correlation and score variance for different methods. (***a***) RGEN-seq *vs* SITE-seq: score correlations between shared cut sites from experiments performed on the same cellular source of genomic DNA (HEK293 gDNA) but at four different Cas9 RGEN concentrations using sgRNAs against FANCF, EMX1, and CD151 gene targets. Correlation between four RGEN concentrations was calculated using Pearson’s correlation coefficient (***b***) Comparison of score variance (log scale) between four methods. Off-target sites are grouped according to number of mismatches (the 3-mismatch group includes sites with 0-3 mismatches).

Though it is obvious RGEN-seq, SITE-seq, and CHANGE-seq are in the same league in regard to their sensitivity, which is substantially higher than that of the remaining two methods, they are not comprehensive, as it is apparent from method comparisons in the intersect and distinct modes (Fig. 4). Indeed, in the three aforementioned methods using FANCF-specific sgRNA, 88 out of 221, 83 out of 223, and 87 out of 164 off-target sites are unique to RGEN-seq, SITE-seq, and CHANGE-seq, respectively. Overall, the percentage of unique off-targets in RGEN-seq, SITE-seq, and CHANGE-seq varies from 25% to 67% for different sgRNAs (Fig. 4*a,b*). It is also true across different method combinations, for example, RGEN-seq and SITE-seq share 116 (FANCF panel) and 457 (VEGFAs1 panel) sites, from which 54 and 96, respectively, are unique for these two methods (Fig. 4*a*). However, all compared methods are much more consistent in determining off-target sites with 1-3 mismatches (Supplementary Fig. s3*a,b*). The latter data also reveal RGEN-seq produces the most output of off-target sites with 6-7 mismatches. It is also worthwhile to note that the number of distinct off-target sites recovered with different software pipelines can vary substantially (Supplementary Fig. s4).

Besides the analysis of intersecting and distinct off-target sites in individual methods and/or methods combinations, we examined the performance of scores assigned to off-targets by BLENDER^22^, which is a sum of the read ends in a 10-bp window centered on the cut site. Figure 5A compares RGEN-seq and SITE-seq in regard with the score correlation between off-target sites identified at different Cas9 RGEN concentrations in the range of 16-1024 nM. For each method only intersecting cut sites at all four concentrations were used for correlation calculation. Overall, RGEN-seq demonstrates a better score correlation, especially in the experiments performed at the lowest and highest RGEN concentrations. Of the four methods producing the highest number of off-targets, RGEN-seq shows the most monotonic change in scores, closely following with SITE-seq, which is reflected by the lowest score variance across off-target sites grouped by number of mismatches (Fig. 5*b*). It is interesting that if we stratify off-target sites by supporting score, the target sites in the CHANGE-seq and CIRCLE-seq methods always occur on top with substantial gap in score between the target and off-target cuts for seven out of eight tested sgRNAs (Supplementary Table s2); however, this is rarely the case for RGEN-seq or SITE-seq. The crucial difference between RGEN-seq and SITE-seq on one side and CHANGE-seq and CIRCLE-seq on the other side is that in the latter two methods RGENs operate on closed circular DNA rather than on linear DNA, and the used RGEN-to-DNA ratio is much higher (∼ 30 fold) than one used in RGEN-seq and SITE-seq.

## DISCUSSION

The method described here, RGEN-Seq, allows selective biochemical genome-wide mapping of Cas9 cleavage sites within genomic DNA. In terms of the overall workflow RGEN-seq is close to DIGENOME-seq, which is by far the simplest existing biochemical off-target site detection technique. Indeed, RGEN-seq is unbiased like DIGENOME, and adds just one extra modified adapter ligation step to the otherwise standard NGS library workflow but that step makes a huge difference in terms of the sensitivity of off-targets site detection. In regards to the latter, it rivals the sensitivity of SITE-seq, CHANGE-seq, or CIRCLE-seq but it employs fewer biochemical steps, doesn’t require DNA circularization, nuclease treatment, biotin-streptavidin enrichment and, most importantly, it doesn’t use PCR amplification during the library preparation. Perhaps, RGEN-seq’s PCR-free workflow might account for its resulting monotonic cleavage score distribution between off-target sites as well as its robust off-target cut site score correlation in the wide range of RGEN concentrations. Because of the amplification-free nature of RGEN-seq, each read contributing to off-target site scores originates from an individual DNA molecule that has been cleaved by Cas9. We therefore hypothesize that scores produced by RGEN-seq could be more relevantly used as a proxy to quantify the affinity of Cas9:sgRNA complexes to adjacent DNA sites and cleavage efficiency compared to those in PCR-based methods.

Despite the high sensitivity of RGEN-seq and other compared methods, the recovered supersets of potential off-target cleavage sites for a nuclease are not comprehensive. While relatively low number of reads (∼ 2 Million) used for comparisons can partly account for that, there are many other unaccounted factors in datasets generated in different laboratories that could be involved in or at least can contribute significantly to the observed method-specific off-target sites, so it would be premature to generalize the results of this limited comparison. Different gDNA preparations (although from the same cell lines), different sources of Cas9 nucleases with unknown specific activity, different sgRNAs – all these factors can introduce substantial variability in the supersets of off-target sites generated by any method. To wit, all compared methods demonstrate excellent reproducibility between technical replicates if performed in the same laboratory with the same reagents^25–28^.

In terms of the identified off-target sites, RGEN-seq demonstrates the highest concordance with SITE-seq, while CHANGE-seq and CIRCLE-seq form another concordant pair. Such method grouping is not surprising, considering that both RGEN-seq and SITE-seq share several common steps including the same conditions for the cleavage of long linear DNA by Cas9. By contrast, in CIRCLE-seq and CHANGE-seq closed circular DNA of different sizes gets cleaved by Cas9. Depending on circularization conditions such as, for example, the ∼20°C temperature difference between the circularization and Cas9 cleavage reactions^26,27^, closed circular molecules can be under the helix stabilizing torsional stress and even be partially positively supercoiled at the cleavage conditions, depending on the DNA size^41–43^. Such torsional stress and associated positive supercoiling energy can impose a significant barrier for the Cas9:sgRNA-medited R-loop formation^44,45^, resulting in negative selection of off-target sites with lower affinity to Cas9:sgRNA complexes. Even more, Cas9 must engage in multiple aborted rounds of binding and R-loop formation before successfully cleaving the DNA site, but such random non-productive binding can further increase the superhelical density affecting site occupancies by an anti-cooperative mechanism^44^. Perhaps, those phenomena can account for the much higher (∼ 30X higher) Cas9:sgRNA to DNA ratio required in CHANGE-seq and CIRCLE-seq than in RGEN-seq and SITE-seq to achieve a comparable level of off-target site detection sensitivity. It also could explain the very high score of target sites compared to any off-target sites in the CHANGE-seq and CIRCLE-seq, as well as higher score variance in these methods in comparison with RGEN-seq and SITE-seq.

In our work we predominantly applied the BLENDER pipeline for detecting off-target sites in all methods but DIGENOME-seq, and in a few cases for comparison we used SITE-seq and DIGENOME pipelines. All these pipelines rely on counting reads with identical 5’ ends around cut sites but implement it differently, so not surprisingly different algorithms produce both intersecting and distinct off-targets with the same dataset (Supplementary Fig. s4). While comprehensive evaluation of the algorithms is beyond the scope of this study, we noticed that often off-target cleavage sites identified as unique by a given pipeline for a given method happen to not be unique after visual inspection of the alignment. Thus, the off-target catching software may contribute to the observed differences in the supersets of cleavage sites for a given sgRNA and more accurate predictive algorithms for Cas9 off-target detection is required.

In conclusion, *in vitro* detection of potential off-target double stranded breaks produced by gene editors is an essential part of their characterization and simple, sensitive, scalable, easily automatable, and cost effective off-target detection methods are very important for an initial screens to characterize CRISPR-Cas nucleases. RGEN-seq is well positioned to fill this niche and it can be easily implemented in both a small academic laboratory and in a high-throughput automated facility.

## Methods

### Nucleic acids and enzymes

Primers and gRNAs used in this work were purchased from Integrated DNA Technologies (IDT) and their sequences are available in Supplementary Table s0. For experiments with CHO-K1 DNA custom synthesized 36-mer crRNAs were annealed to universal 67-mer tracrRNA to form sgRNA; for human genomic DNA full size 100 b sgRNAs were synthesized. Genomic CHO-K1 DNA from the AD49GZ clone was isolated as previously described^30^; human Hek293 DNA was purchased from Genscript. *S. pyogenes* Cas9, calf intestinal phosphatase (CIP), terminal transferase (TdT), Klenow Fragdment (3’→5’ exo-), Quick T4 DNA ligase were from New England Biolabs (NEB); RNase cocktail™ was from Thermo Fisher Scientific; Proteinase K was from Qiagen.

### RGEN-seq library preparation and sequencing

When indicated, genomic DNA was pre-treated with CIP and TdT with ddATP according to the manufacturer’s protocol to prevent adapter ligation to pre-existing random double-stranded breaks (DSBs). RGEN complexes were assembled by mixing *S. pyogenes* Cas9 with sgRNA (ratio 1:1.5) at room temperature for 10 minutes. In multiplexing experiments, either sgRNAs were mixed in equimolar amounts prior to mixing with Cas9, or, alternatively, individual sgRNA were mixed with Cas9 and then the resulting complexes were combined in equal proportions. For each reaction, 1-4 μg of genomic DNA (gDNA) was mixed with a corresponding RGEN at 16-1024 nM and incubated at 37°C for 1 hour; then the reaction was stopped by heating to 65°C for 10 minutes. The terminated RGEN reactions were treated with RNase cocktail and Proteinase K, followed by purification of cleaved gDNA with Beckman SPRIselect paramagnetic beads according to the manufacturer’s instructions. Purified cleaved gDNA was end repaired and A-tailed with Klenow Fragment (3’→5’ exo-), followed by ligating DNA with indexed p7L-adapter and purification with SPRIselect beads to remove free adapter. The resulting DNA (500 ng) was concurrently fragmented, end repaired, and A-tailed using NEBNext Ultra II FS DNA Module (NEB); fragmented DNA then was ligated with p5L adapter using NEBNext Ultra II Ligation Module (NEB). After the SPRIselect purification, completed libraries were assessed on Bioanalyzer for fragment sizes, quantified by qPCR and loaded on an Illumina sequencer. Indexed RGEN-seq libraries were sequenced on an iSeq100, NextSeq550, or NS2000 using paired-end sequencing with R1×26 and R2×151 cycles. A detailed user protocol for RGEN-seq library construction is provided (Supplementary Protocol)

### RGEN-seq analysis and off-target sites identification

R2 reads were mapped to the reference human genome (hg38) using bwa mem with default settings, then converted to bam format, indexed, and sorted using samtools. The resulting indexed and sorted bam file was used as an input for BLENDER, SITE-seq, or DIGENOME-seq off-target calling pipelines. For RGEN-seq comparison with other biochemical methods, corresponding sequencing reads were downloaded from the NCBI’s Sequence Read Archive (SRA) and processed in the same way as RGEN-seq data. The genomic coordinates in the DIGENOME-seq, identified off-target sites based on the hg19 version of human genome, were converted to the hg38 version using the NCBI’s Remap tool. The benchmark study focused on the traditional CRISPR SpCas9 system and utilized gRNAs with 20 nucleotides (nt) followed by NGG-PAM. Statistical analyses were performed in R and results were plotted using R’s ggplot2, UpSet, ComplexHeatmap, and PerformanceAnalytics packages. For visual analysis of cut sites, Integrated Genomic Viewer^46^ and NCBI’s Genome Workbench^47^ were used.

## Supporting information

Suppelentary information

Suppementary RGEN-seq protocol

Suppelementary Table s3

Suppelentary Table s0

## Data Availability

High-throughput sequencing reads will be deposited in the NCBI Sequence Read Archive database after publication.

## Author Contributions

Wet lab experiments were performed by AK, JO, and AS. AS, BR, and AK performed informatic analyses. AS wrote an initial draft of the manuscript and all authors contributed to the writing of the final version of the manuscript.

## Notes

### Competing Interest Statement

The authors have declared no competing interest.

