## Supplementary material for "RGEN-SEQ FOR HIGHLY SENSITIVE AMPLIFICATION-FREE SCREEN OF OFF-TARGET SITES OF GENE EDITORS": Suppelentary information

Alexander Kuzin, Brendan Redler, Jaya Onuska, Alexei Slesarev*
BioReliance Corp., 14920 Broschart Road, Rockville, MD 20850, USA

**Supplementary Table s0**. List of sequences of sgRNAs and NGS adapters.
See attached file.

**Supplementary Table s1**. Laboratory components used for RGEN-seq optimization
See attached file.

**Supplementary Table s2**. List of off-target sites used in Figures 3 and 4.
See attached file

**Supplementary Table s3**. List of off-target sites used in Figure 5.
See attached file.

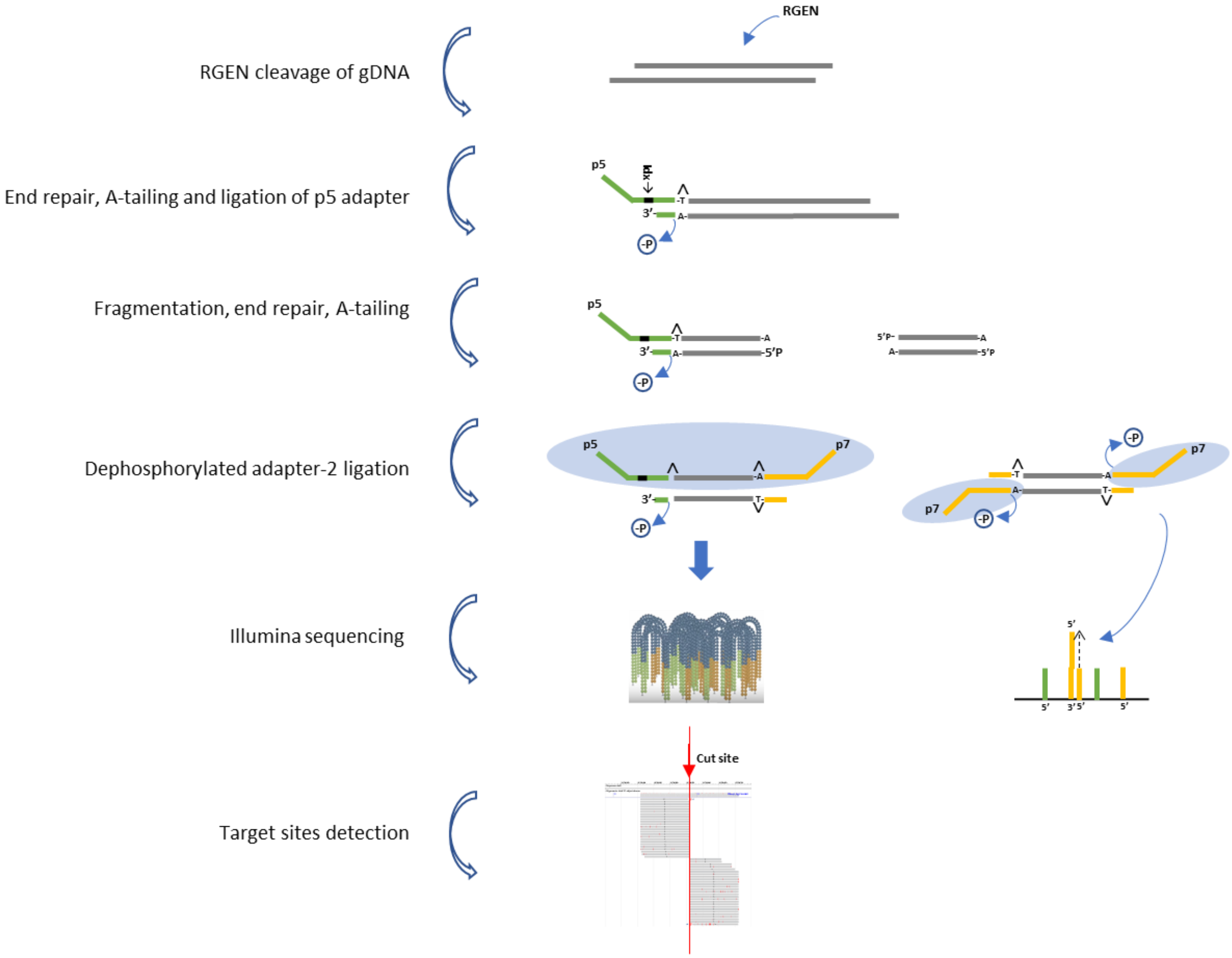


**Supplementary Figure s1**. An alternative version of RGEN-seq workflow. Schematic workflow illustrating preparation of the RGEN-seq library compatible with Illumina single-read sequencing. The p5L-adapter, which is an Illumina Y-type index adapter (orange), is missing the p7 site; the p7L-adapter (green), which is a truncated Illumina Y-adapter, lacks the p5 site. Ligation of the latter adapter to both sides of free fragmented DNA prevents bridge amplification of “background” library molecules, though free p7L-adapter still can anneal the flowcell-bound oligonucleotides. Target and off-targets cut sites are detected by spotting the characteristic read alignment pattern around cleavage sites.

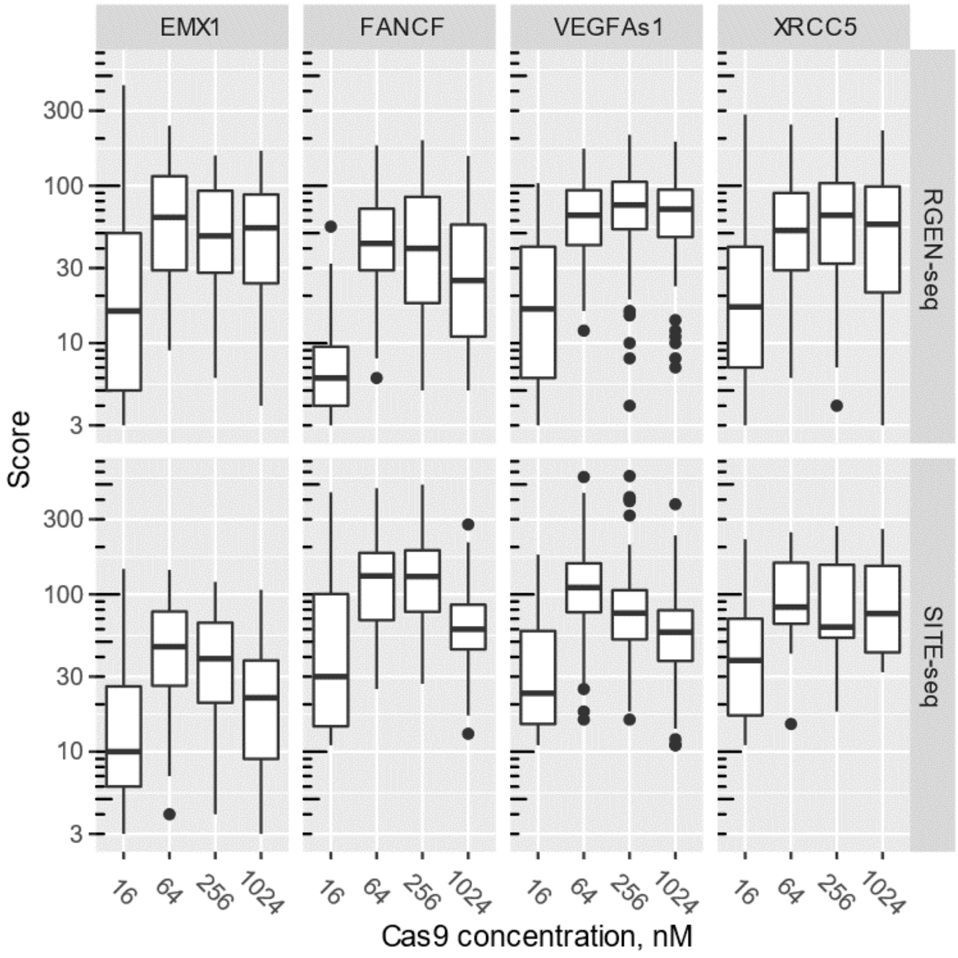


**Supplementary Figure s2.** Boxplot representation of the cleavage sites score distributions as a function of RGEN concentration. Two methods, RGEN-seq and SITE-seq, are compared using four different targets.

*a*

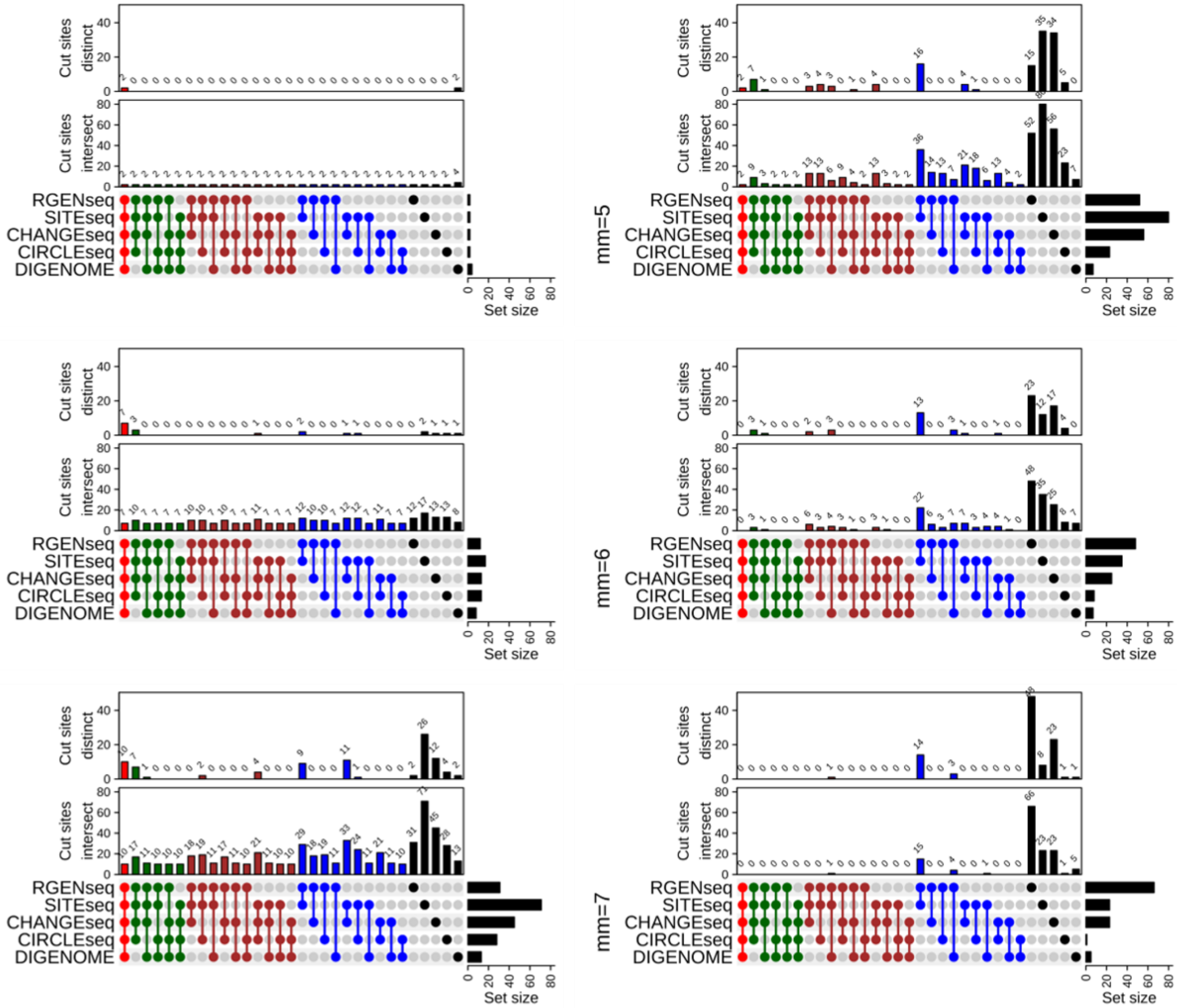

mm=[0,2]
mm=3
mm=4
mm=5
mm=6
mm=7
Cut sites distinct
Cut sites intersect
RGENseq
SITEseq
CHANGEseq
CIRCLEseq
DIGENOME
Set size

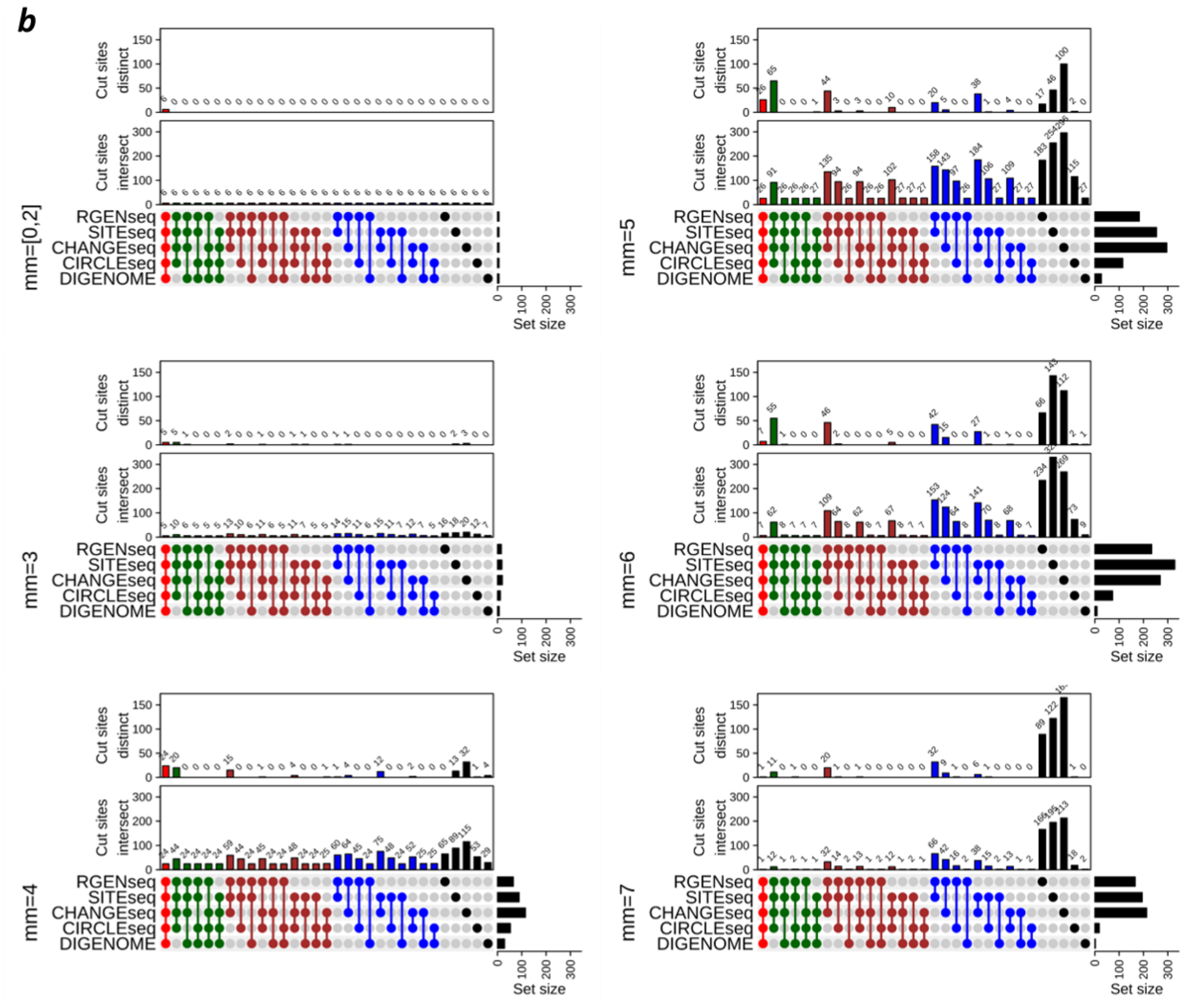


**Supplementary Figure s3. Comparison of intersecting and distinct Cas9 cut sites grouped by number of mismatches across different biochemical methods**. UpSet plots of intersecting (lower plot) and distinct (upper plot) off-target sites of sgRNAs against FANCF (***a***) and VEGFAs1 (***b***) genes produced by five methods.

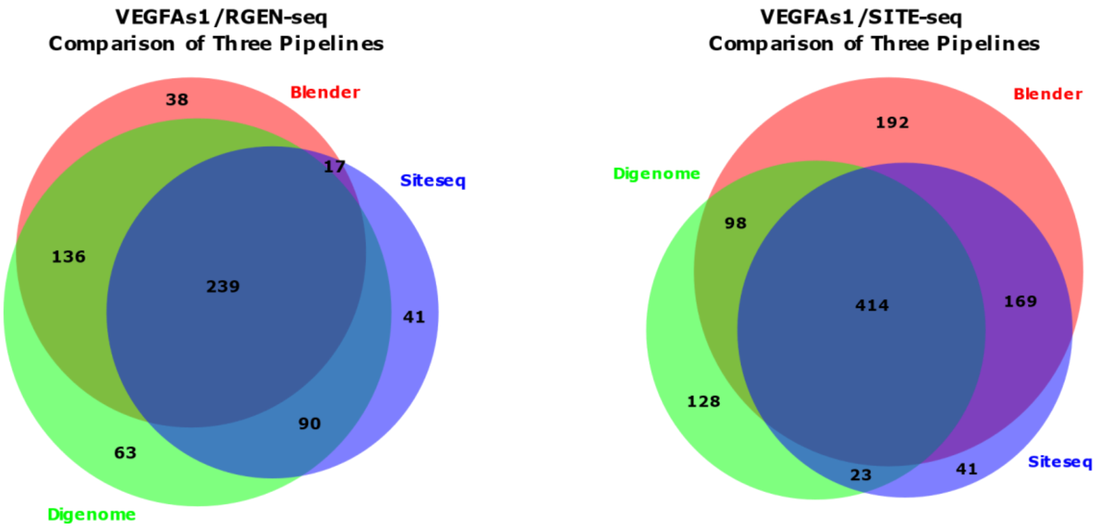


**Supplementary Figure s4. Comparison of three different off-target calling pipelines**. Venn diagrams showing the number of overlapping sites identified by three different pipelines. Off-target sites of VEGFAs1 sgRNA captured by RGEN-seq and SITE-seq were analyzed using DIGENOME[25], BLENDER[22], and SITE-seq[28] off-target calling software.

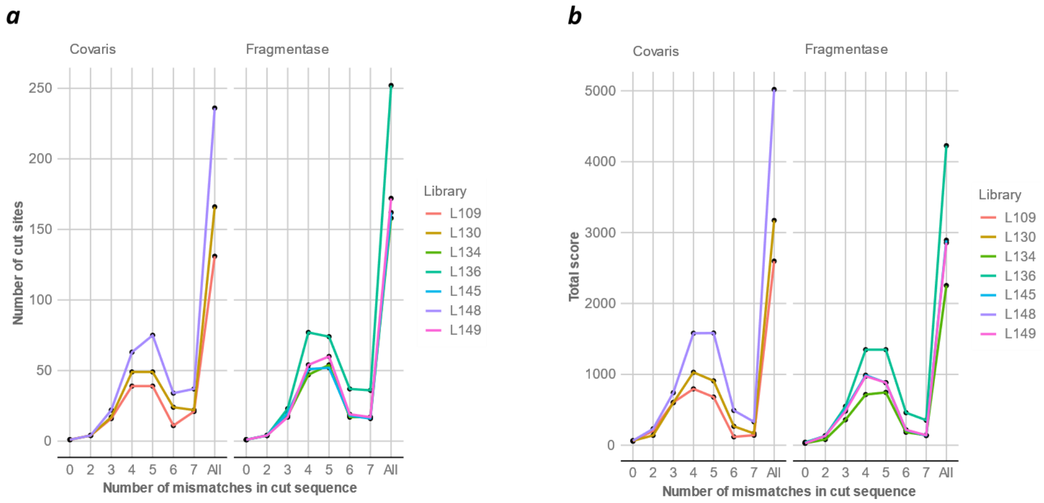


**Supplementary Figure s5. Effect of different DNA fragmentation methods on recovery of off-target cut sites in RGEN-seq**. Two DNA fragmentation methods are compared, ultrasound shearing using a Covaris instrument and an enzymatic fragmentation with Fragmentase. (***a***), Shown are numbers of detected off-target cut sites grouped by number of mismatches between the guide sequence (EMX1 sgRNA) and cut site sequences. In (***b***) total scores for individual mismatch groups of off-target cut sites are shown.

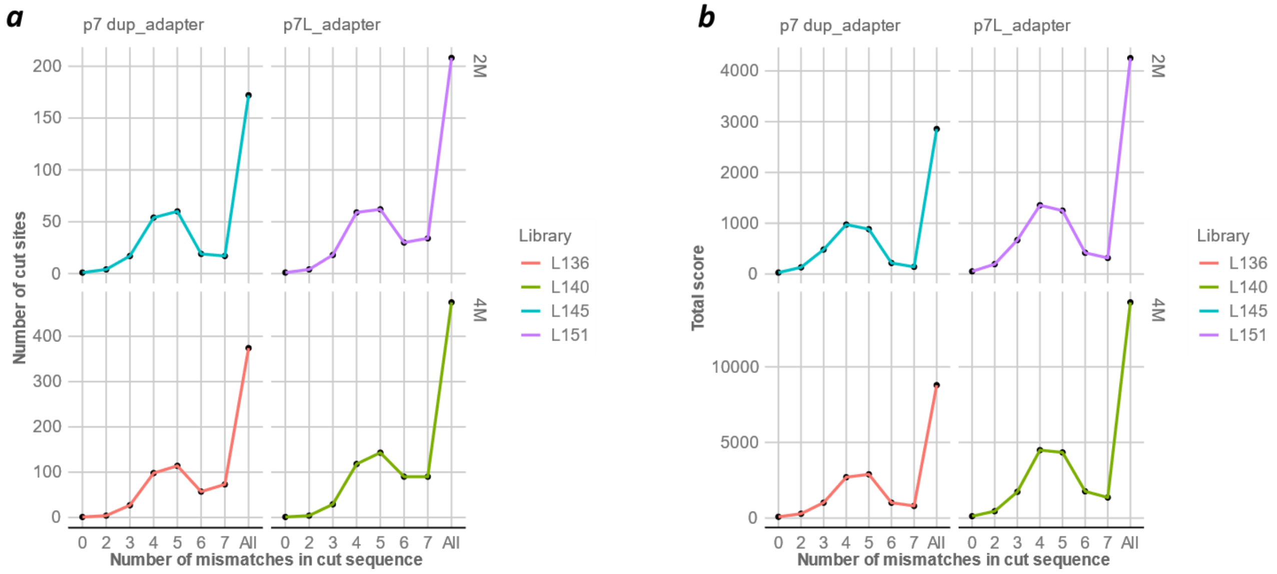


**Supplementary Figure s6. Effect of first adapter structure on recovery of off-target sites in RGEN-seq**. The p7 duplex adapter is compared with the partial dupex p7L using two metrics: numbers of off-target cut sites grouped by number of mismatches (***a***), and total scores for individual mismatch groups of off-target cut sites (***b***)**.** Two million (*top panels*) and four million (*bottom panels*) reads were used for detecting off-target cut sites produced with EMX1 RGEN.

***a***

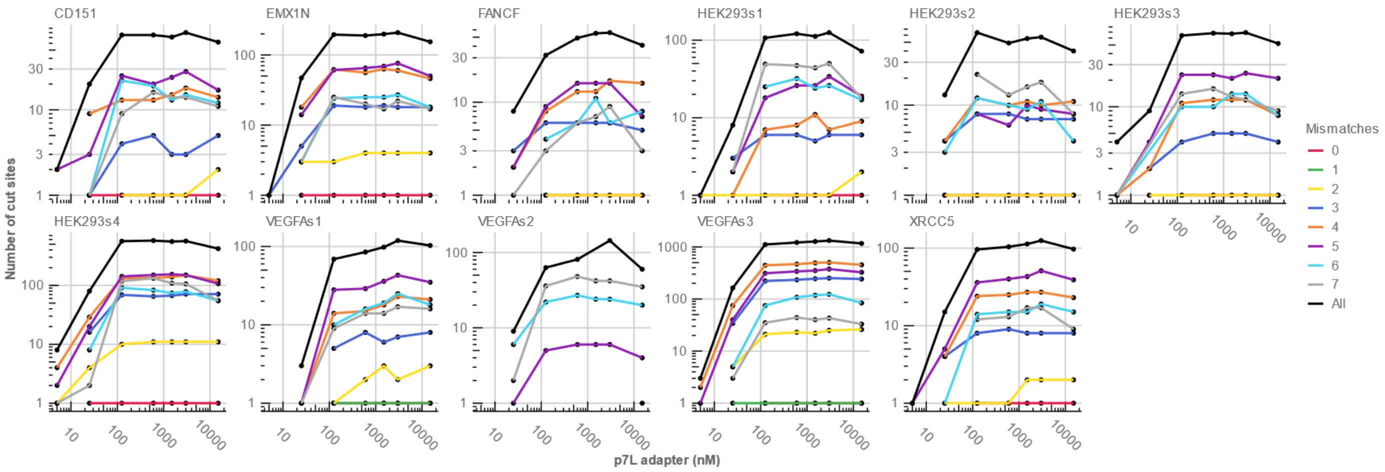


***b***

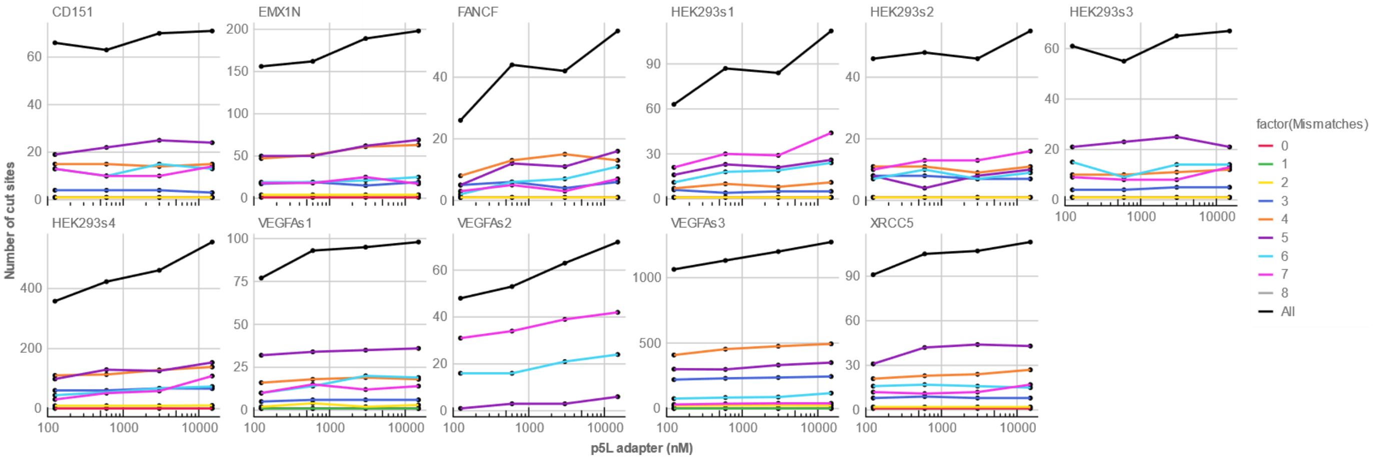


**Supplementary Figure s7. Effect of first adapter concentrations on number of detected off-target cut sites in RGEN-seq**. Shown numbers of off-target cut sites are for the first p7L adapter (**a**) and for the second p5L adapter (***b***). Color coded curves correspond to the specified number of mismatches between guide sequences and cut site sequences.

### SUPPLEMENTARY NOTE

**RGEN-seq optimization**.

We directed our major optimization efforts to the reduction of background reads. Such reads are produced by ligating the first adapter to pre-existing DSBs in genomic DNA and to DSBs, which are generated during purification steps. We reduced the number of purification steps from seven in the initial RGEN-seq version to just four in the final version by combining several different reaction steps into a single reaction. Figure s5 demonstrates that substitution of ultrasound shearing with enzymatic fragmentation eliminated one purification step without sacrificing the overall efficiency of RGEN-seq. One more purification step was eliminated by using Klenow Fragment (3'-->5' exo-) DNA polymerase with dNTPs to combine the end repair (ER) and dA-tailing steps into one after the RGEN cleavage step; initially we used Illumina's library ER and dA-tailing procedure, which required an intermediate sample purification.

Further optimization was performed to improve the first adapter ligation step. We initially used the p7 duplex adapter obtained by annealing the p7_index oligonucleotide to its complement variant but this p7 duplex was then relaced with the p7L adapter. The latter is easier to remove from the ligation reaction due to its smaller size, which is important as any carry-over of residual adapter 1 into the next step can increase the background reads. This replacement resulted in somewhat higher sensitivity of RGEN-seq (Fig. s6).

Next, we evaluated the effect of different concentrations of adapter 1 and adapter 2 on the number of recovered off-target cut sites; based on the results presented in Figure s7 we determined the optimal concentration of the p7L and p5L adapters as 1.5 µM and 15 µM, respectively.

Finally, we developed the continuous high-throughput RGEN-seq method for the Perkin Elmer Sciclone G3 NGSx liquid handling workstation allowing walkaway RGEN-seq library preparation. The automated method consists of optimized enzymatic reactions followed by magnetic bead-based purification steps, with appropriate liquid handling parameters to ensure efficient reactions and pre-digested gDNA integrity.
