## Supplementary material for "RGEN-SEQ FOR HIGHLY SENSITIVE AMPLIFICATION-FREE SCREEN OF OFF-TARGET SITES OF GENE EDITORS": Suppementary RGEN-seq protocol

Alexander Kuzin, Brendan Redler, Jaya Onuska, Alexei Slesarev*
BioReliance Corp., 14920 Broschart Road, Rockville, MD 20850, USA

### Reagents

1. HEK293 genomic DNA (gDNA) (Genscript, M00094)
2. 10X CutSmart Buffer (NEB, B7204)
3. RT-PCR Grade Water (Invitrogen, AM9935)
4. Nuclease Free Duplex Buffer (IDT, 11-01-03-01)
5. *S. pyogenes* Cas9 (NEB, M0386)
6. sgRNA of Interest (IDT, custom)
7. RNase Cocktail (ThermoFisher, AM2286)
8. Proteinase K (QIAGEN, 19131)
9. SPRISelect Reagent Kit (Beckman CoµLter, B23317)
10. Qubit HS DNA Kit (Invitrogen, Q32854)
11. (3’->5’ exo$^-$) Klenow Fragment (NEB, M0212)
12. 10mM dNTP (NEB, N0447)
13. NEBNext Quick T4 DNA Ligase (NEB, E6056)
14. ATP (NEB, P0756)
15. NEBNext MLtra II FS DNA Module (NEB, E7810)
16. NEBNext MLtra II Ligation Module (NEB, E7595)
17. Zymo Research Select-a-Size DNA Clean and Concentrator columns (Zymo Research, D4080)
18. Agilent DNA 1000 reagents kit (Agilent, 5067-1504)
19. KAPA Library Quant Kit KK4824 (Roche, 07960140001)
20. Ethyl Alcohol, pure (Sigma-Aldrich, E7023)
21. Oligonucleotides (IDT, custom)
22. 10mM ddATP (Sigma, GE27-2051-01)
23. Quick CIP (NEB, M0525)
24. Terminal Transferase (NEB, M0315)

| | |
|---|---|
| rc_p7 | GCTCTTCCGATC*T |
| p7_index | /5’Phos/GATCGGAAGAGCACACGTCTGAACTCCAGTCACNNNNNNCGATGTATCTCGTATGCCGTCTTCTGCTTG/3’AmMO/ |
| rc_p5 | GATCGGAAGAGCGTCGTG |
| p5_index | AATGATACGGCGACCACCGAGATCTACACNNNNNNNNACACTCTTTCCCTACACGACGCTCTTCCGATC*T |

### 0. Blocking gDNA fragment ends to minimize non-specific adapter ligation (optional step; it is recommended if genomic DNA is degraded)

1. Remove reagents from -30°C freezer, allow to thaw on ice.
2. Dilute 3μg gDNA in 38μL 1x TE by gentle flicking the tube.
3. Assemble the blocking gDNA reaction by gentle flicking the tube:

| | |
|---|---|
| gDNA | 38.0 μL |
| 10x NEB CutSmart buffer | 5.0 μL |
| 10mM ddATP | 0.5 μL |
| Quick CIP enzyme | 2.0 μL |
| Terminal Transferase | 4.5 μL |
| Total | 50.0 μL |

4. Incubate the reaction at 37ºC for 60 minutes, then at 75ºC for 20 minutes.

### 0.1 SPRIselect gDNA cleanup for step 0 (skip it if no blocking reaction was performed)

1. Add 50 μL of TE to the gDNA sample and mix gently by flicking the tube.
2. Add 80 μL of resuspended SPRISelect beads to the sample. Mix gently but thoroughly.
3. Incubate the mixture for 5 min at room temperature (RT).
4. Pellet the reaction on a magnetic rack, allow beads to pellet for 2 minutes (until the supernatant is clear and colorless).
5. Pipette off the supernatant.
6. Keep the tube on a magnetic rack and wash the beads with 350 μL of 85% ethanol without disturbing the pellet. Remove supernatant after 30 seconds.
7. Dry the pellet for 30-60 seconds, but do not over-dry as it will resμLt in significant loss of gDNA.
8. Remove the tube from the magnetic rack and resuspend pellet in 39 μL of TE by gently mixing. Incubate for 10 minutes for eluting DNA into solution.
9. Pellet the eluate on the magnetic rack. Keep the tube on the magnet for 2 minutes (until the eluate is clear and colorless).
10. Remove and retain 37 μL of eluted gDNA

### A. gDNA Digestion by RGEN Complex

1. Remove reagents from -30°C freezer, allow to thaw on ice.
2. Prepare 50μM 100 b sgRNA. Heat to 95ºC for 3 minutes and allow to slowly cool to room temperature.
3. Dilute sgRNA(s) and Cas9 nuclease to needed concentrations of the RGEN experiment in 1x NEB CutSmart buffer.
4. Prepare the RGEN complex by mixing diluted sgRNA(s) and Cas9 enzyme with sgRNA(s): Cas9 ratio of 1.5:1 in 10μL 1x NEB CutSmart buffer.
5. Incubate the RGEN complex at RT for 10 minutes.

6. Dilute 1-4 μg gDNA in 40μL 1x NEB CutSmart buffer by gently mixing.
7. Add the RGEN complex to diluted gDNA and incubate at 37°C for 60 minutes.
8. Heat up to 65°C for 10 minutes to terminate the reaction.
9. Add 0.5 μL of RNase Cocktail and incubate the reaction at 37°C for 30 minutes.
10. Add 2 μL of Proteinase K and incubate the reaction at 50°C for 30 minutes.

### B. gDNA cleanup 1

1. Add 47.5 μL of TE to the final volume (52.5 μL) from the previous step.
2. Resuspend the SPRISelect beads by vortexing for 1 minute.
3. Add 70 μL of resuspended SPRISelect beads to the sample. Mix thoroughly but gently. Incubate for 5 minutes at RT.
4. Prepare 85% ethanol.
5. Pellet the reaction on the magnetic rack for 2 minutes (until the supernatant is clear and colorless) and pipette off the supernatant.
6. Keep the tube on magnet and wash the beads with 350 μL of 85% ethanol without disturbing the pellet for 30 seconds. Remove all ethanol and discard.
7. Dry the pellet for 30-60 seconds, but do not over-dry. If pellet is overdried gDNA may be lost.
8. Remove the tube from the magnetic rack and resuspend pellet in 33 μL of TE by gentle mixing. Incubate for 10 minutes for eluting DNA into solution.
9. Pellet the eluate on a magnetic rack for 2 minutes (until the eluate is clear and colorless).
10. Remove and retain 31 μL of eluted gDNA.

### C. dA-tailing and the p7L adapter ligation

1. Remove reagents from -30°C freezer, allow to thaw on ice.
2. Prepare 1.5μM p7L adapter as follows:

| | |
|---|---|
| p7_index (100 uM) | 1 μL |
| rcp7 (100 uM) | 1 μL |
| IDT Duplex buffer | 64.5 μL |
| Total | 66.5 μL |

3. Incubate p7L adapter at 95°C for 3 minutes and then allow it to slowly cool to RT.
4. Assemble the dA-tailing reaction:

| | |
|---|---|
| RGEN-treated gDNA from step B | 30.25 μL |
| 10x NEB CutSmart buffer | 3.75 μL |
| 10mM dNTP | 0.5 μL |
| (3’->5’ exo$^{-}$) Klenow Fragment | 3 μL |
| Total | 37.5 μL |

5. Incubate the reaction at 37°C for 30 minutes and heat up to 75°C for 20 minutes to terminate dA-tailing reaction.
6. Assemble the first ligation reaction:

| | |
|---|---|
| dA-tailed gDNA | 37.5 μL |
| 10x T4 DNA ligation buffer | 1.5 μL |

| | |
|---|---|
| 10mM ATP | 3.5 μL |
| 1.5μM First Adapter | 2.5 μL |
| NEBNext Quick T4 DNA ligase | 5 μL |
| Total | 50 μL |

7. Incubate the reaction at RT for 30 minutes.

**D. gDNA cleanup 2**

1. Add 50 μL of TE to the final volume (50 μL) from the previous step.
2. Resuspend the SPRISelect beads by vortexing for 1 minute.
3. Add 70 μL of resuspended SPRISelect beads to the sample. Mix thoroughly but gently. Incubate for 5 minutes at RT.
4. Prepare 85% ethanol.
5. Pellet the reaction on the magnetic rack for 2 minutes (until the supernatant is clear and colorless) and pipette off the supernatant.
6. Keep the tube on magnet and wash the beads with 350 μL of 85% ethanol without disturbing the pellet for 30 seconds. Remove all ethanol and discard.
7. Dry the pellet for 30-60 seconds, but do not over-dry. If pellet is overdried gDNA may be lost.
8. Remove the tube from the magnetic rack and resuspend pellet in 32 μL of TE by mixing. Incubate for 10 minutes.
9. Place the eluate on a magnetic rack. Keep the tube on the magnet for 2 minutes (until the eluate is clear and colorless).
10. Remove and retain 30 μL of eluted gDNA.

**E. gDNA fragmentation and end-repair/dA-tailing**

1. Remove reagents from -30°C freezer, allow to thaw on ice.
2. Assemble the dA-tailing reaction as follows:

| | |
|---|---|
| 150-500ng gDNA | 26 μL |
| NEBNext MLtra II FS Reaction buffer | 7 μL |
| NEBNext MLtra II FS Enzyme Mix | 2 μL |
| Total | 35 μL |

3. Incubate the reaction at 37°C for 12 minutes and then at 65°C for 30 minutes.

**F. p5L adapter ligation**

1. Prepare 15μM p5L adapter as follows:

| | |
|---|---|
| p5_index (100 uM) | 1 μL |
| rc_p5 (100 uM) | 1 μL |
| IDT Duplex buffer | 4.6 μL |
| Total | 6.6 μL |

2. Incubate p5L adapter mix at 95°C for 3 minutes and then allow it to slowly cool to RT.

3. Assemble the second ligation reaction as follows:

| | |
|---|---|
| End-repaired DNA from sep E | 35 µL |
| 15µM p5L adapter | 2.5 µL |
| NEBNext MLtra II Ligation Master Mix | 30 µL |
| NEBNext Ligation Enhancer | 1 µL |
| Total | 68.5 µL |

4. Incubate the reaction at RT for 20 minutes.

**G. gDNA cleanup 3**

1. Resuspend the SPRISelect beads by vortexing for 1 minute.
2. Add 31.5 µL of TE. Mix.
3. Add 70 µL of resuspended SPRISelect beds. Mix. Incubate for 5 minutes.
4. Prepare 85% ethanol.
5. Pellet the reaction on a magnetic rack. Keep the tube on the magnet for 2 minutes (until the supernatant is clear and colorless) and pipette off the supernatant.
6. Keep the tube on magnet and wash the beads with 350 µL of 85% ethanol without disturbing the pellet for 30 seconds. Remove all 85% ethanol drops using a pipette tip and discard.
8. Dry the pellet for 30-60 seconds, but do not dry the pellet completely.
9. Remove the tube from the magnetic rack and resuspend pellet in 25 µL of TE by mixing. Incubate for 10 minutes.
10. Pellet the eluate on a magnetic rack. Keep the tube on the magnet for 2 minutes (until the eluate is clear and colorless).
11. Transfer and retain 23 µL of made library into the 1.5mL Eppendorf DNA LoBind tube.

**H. Library quality control, quantification, and sequencing**

1. Analyze library size on an Agilent Bioanalyzer using a DNA 1000 chip.
2. Quantify library using qPCR.
3. Perform paired-end sequencing on an Illumina sequencer. Use 25 cycles for R1 reads and 100-300 cycles (depending on a sequencer used) for R2 reads.

### ALTERNATIVE PROTOCOL OF DNA SHEARING USING COVARIS S220 ULTRASONICATOR

Perform steps A-D as in the previous protocol

**E. DNA shearing**

Consumable: Covaris microtube AFA Fiber Crimp-Cap 6x16mm (Covaris, 520052)

1. Add 32 µL of TE to the DNA sample from the step D and transfer the sample into the Covaris microTUBE.
2. Perform DNA shearing in Covaris S220 (set up parameters: 175W of peak incident power; 5% of duty factor; 200 cycles per burst; 75 seconds of treatment time).
3. Transfer the sample into a 1.5mL Eppendorf DNA LoBind tube.
4. Adjust the sample volume to 100 µL with TE buffer.

**F. DNA cleanup 3**

Reagent for DNA cleanup 3: Zymo Research Select-a-Size DNA Clean and Concentrator columns (Zymo Research, D4080).

1. Prepare Zymo fresh Select-a-Size DNA binding solution by adding 0.5 mL of Zymo Select-a-Size DNA Binding Buffer to 70 µL of 100% ethanol and mix thoroughly.
2. Add 100 µL of DNA sample to Zymo Select-a-Size DNA binding solution from step 1 and mix thoroughly.
3. Apply the mixture onto the Zymo-Spin IC-S column in a collection tube. Centrifuge at 11000 x g for 30 seconds. Discard the flowthrough.
4. Add 700 µL of Zymo DNA Wash buffer to the column. Centrifuge at 11000 x g for 30 seconds. Discard the flow-through.
5. Add 200 µL of Zymo DNA Wash buffer to the column. Centrifuge at 11000 x g for 60 seconds. Discard the collection tube.
6. Transfer the column to a 1.5mL Eppendorf DNA LoBind tube and add 54 µL of Zymo DNA Elution Buffer to the column matrix and incubate for 5 minutes at room temperature. Centrifuge at 11000 x g for 30 seconds to elute the DNA.

**G. End-repair/dA-tailing**

Reagents: NEBNext Ultra II End Repair/dA-Tailing Module (NEB, E7546)

1. Remove reagents from -30°C freezer, allow to thaw on ice.
2. Assemble the end-repair/dA-tailing reaction as follows:

| | |
|---|---|
| Purified fragmented DNA | 50µL |
| NEBNext Ultra II End Prep Reaction Buffer | 7µL |
| NEBNext Ultra II End Prep Enzyme Mix | 3µL |
| Total | 60µL |

3. Incubate the reaction at 20°C for 30 minutes and then at 65°C for 30 minutes.

**H. p5L adapter ligation**

1. Prepare 15µM the second adapter. Mix and dilute tua-s olig and rc-tua_noP7 olig. Heat to 95°C for 3 minutes and allow to slowly cool to room temperature.
2. Assemble the second ligation reaction as follows:

| | |
|---|---|
| Fragmented DNA | 60µL |
| 15µM Second Adapter | 2.5µL |
| NEBNext Ultra II Ligation Master Mix | 30µL |
| NEBNext Ligation Enhancer | 1µL |

| | |
|---|---|
| Total | 93.5µL |

3. Incubate the reaction at RT for 20 minutes.

### I. DNA cleanup 4

Reagent for DNA cleanup 4: Zymo Research Select-a-Size DNA Clean and Concentrator columns (Zymo Research, D4080)

1. Add 6.5 µL of Zymo DNA Elution Buffer to the sample and mix.
2. Prepare Zymo fresh Select-a-Size DNA binding solution by adding 0.5 mL of Zymo Select-a-Size DNA Binding Buffer to 70 µL of 100% ethanol and mix thoroughly.
3. Add 100 µL of DNA sample to Zymo Select-a-Size DNA binding solution from step 1 and mix thoroughly.
3. Apply the mixture onto the Zymo-Spin IC-S column in a collection tube. Centrifuge at 11000 x g for 30 seconds. Discard the flowthrough.
4. Add 700 µL of Zymo DNA Wash buffer to the column. Centrifuge at 11000 x g for 30 seconds. Discard the flow-through.
5. Add 200 µL of Zymo DNA Wash buffer to the column. Centrifuge at 11000 x g for 60 seconds. Discard the collection tube.
4. Transfer the column to a 1.5mL Eppendorf DNA LoBind tube and add 35 µL of 0.1x Zymo DNA Elution Buffer to the column matrix and incubate for 5 minutes at room temperature. Centrifuge at 11000 x g for 30 seconds to elute the DNA.
5. Save the prepared library in the 1.5 mL Eppendorf DNA LoBind tube.

### J. Library quality control, quantification, and sequencing

1. Analyze library size on an Agilent Bioanalyzer using a DNA 1000 chip.
2. Quantify library using qPCR.
3. Perform paired-end sequencing on an Illumina sequencer. Use 25 cycles for R1 reads and 100-300 cycles (depending on a sequencer used) for R2 reads.
